# Target-enriched sequencing enables genomic characterization within diverse microbial populations – a preprint

**DOI:** 10.1101/2025.10.23.684174

**Authors:** Enrique Doster, Lee J. Pinnell, Cory A. Wolfe, Noelle R. Noyes, Robert Valeris-Chacin, William B. Crosby, Michael L. Clawson, Amelia R. Woolums, Paul S. Morley

**Author notes:** Corresponding author: Paul S. Morley.

## Abstract

Characterizing microbial genetic sequences and key variants is critical for understanding pathogen ecology, transmission, and clinical impact. Yet, conventional metagenomic sequencing often yields too few on-target reads to move beyond species-level identification. We developed a target-enriched (TE) metagenomic workflow, including bait design, an optimized TE shotgun protocol, and the VARIANT++ pipeline, to recover and classify reads at a clustered genomic sequence-variant (GSV) level (see Graphical abstract). The computational component clusters reference genomes by average nucleotide identity, builds a GSV database, and integrates Kraken2, Themisto, and mSWEEP to increase call confidence while reducing false positives. Using Mannheimia haemolytica (*Mh*), the primary cause of bovine respiratory disease, we designed 114,375 DNA baits targeting sequences across 70 reference genomes. TE libraries from nasopharyngeal swabs of feedlot cattle achieved >250-fold increases in on-target *Mh* reads (∼2.5% of non-host reads on average) compared with conventional shotgun sequencing, despite using one-quarter the sequencing depth. This variant-level resolution revealed six GSVs; most samples contained at least two, indicating variant mixtures difficult to detect with culture- or shotgun-based surveys. Because the approach leverages available reference sequences, it can be reconfigured for other microbial targets. TE metagenomics paired with genome-similarity clustering provides a scalable approach to variant-level characterization from complex microbial populations.

Graphical abstract
Overview of the components in our three-part workflow.

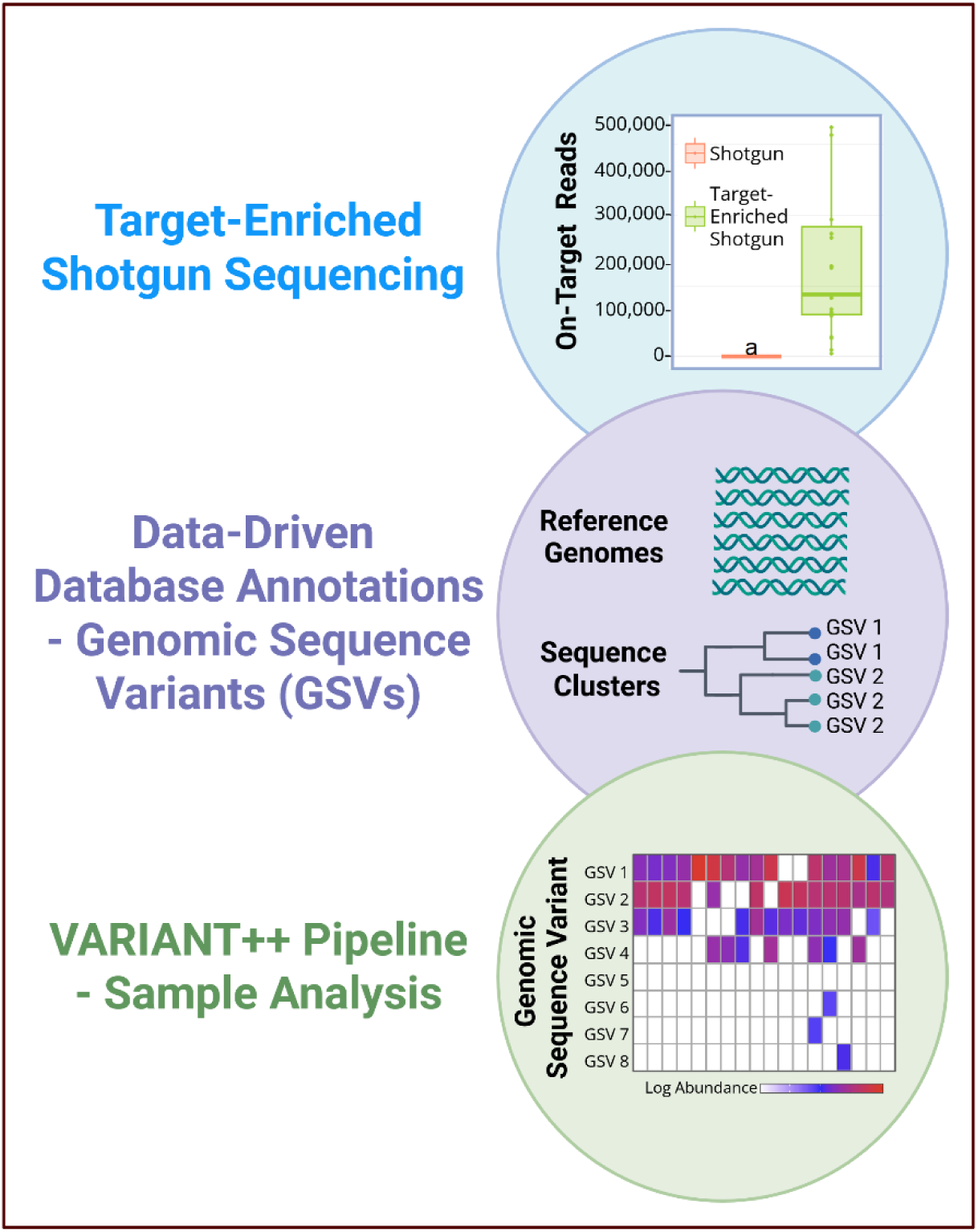

## Main

Characterizing microbial genetic sequences and understanding the significance of key variants within microbial taxa or gene groups of interest is increasingly critical in advancing our understanding of their ecological roles, transmission dynamics, and clinical relevance. However, characterizing any microbial population at the variant level is a formidable challenge due to the lack of a standardized threshold for the genetic variability that defines a “strain”, which results in very inconsistent strain-level classification, even in well-curated databases like NCBI’s RefSeq. Variant-level characterization of microbial targets can provide critical insights into traits such as pathogenicity, antimicrobial susceptibility, host specificity, and ecological fitness, factors that often cannot be inferred from higher-level taxonomic classifications alone^1–4^. Yet, capturing variant-level diversity in a holistic or ecologically relevant context remains exceedingly difficult due to the inherent limitations associated with culture-based methods and culture-independent sequencing approaches, each of which offers only a limited perspective of the underlying microbial landscape.

Culture-based approaches, like whole-genome sequencing (WGS), can yield detailed insight into variant-level physiology, metabolism, and pathogenicity, but they cannot provide information on uncultivable organisms and fail to reflect the broader diversity of microbes relevant to environmental or host-associated ecosystems. Matrix-assisted laser desorption/ionization, time of flight (MALDI TOF) mass spectrometry, for example, has been demonstrated to be a rapid and inexpensive method for distinguishing genotypes^5^, but like WGS, it remains constrained by the need for cultured isolates and offers limited ecological context. Overcoming these constraints requires methods that do not depend on culture, and commonly used approaches like 16S rRNA gene sequencing and shotgun metagenomic sequencing offer solutions for directly analysing microbial communities^6,7^. While both methods have merit depending on the research question, neither can effectively capture variant-level dynamics within microbial populations: 16S rRNA amplicon sequencing suffers from low taxonomic resolution, consistently failing to classify the majority of sequences beyond the rank of genus, while shotgun metagenomic data faces challenges with sensitivity and specificity due to the inclusion of substantial amounts of non-target sequences.

Target-enriched (TE) metagenomic sequencing provides researchers with a method to enrich specific target sequences within a shotgun metagenomic sequencing library and can effectively improve on-target sequencing performance without the need for culture. In addition to targeting important bacterial and viral pathogens^8,9^, TE has been effectively used to enrich antimicrobial resistance genes (ARGs), typically present at extremely low abundance, from diverse microbial populations, resulting in a significant increase in the number of ARGs within metagenomic libraries^10–12^. By increasing the recovery of specific taxa or functional gene groups from complex communities, TE metagenomics offers a scalable and culture-independent strategy for ecological and clinical investigations alike. However, even with tremendous enrichment of on-target reads (i.e. from a few thousand reads, to hundreds of thousands), classifying short-read metagenomic data below the rank of species, particularly in the absence of a standardized definition for what constitutes a “strain”, remains a significant challenge, limiting our ability to produce meaningful information about microbial populations at the genomic variant level.

One clear example of the importance of resolving variant-level structure is *Mannheimia haemolytica* (*Mh*), a respiratory pathogen whose within-species diversity has important implications for disease outcomes. Widely considered the most important pathogen associated with bovine respiratory disease (BRD), a disease estimated to result in losses over a $1 billion USD annually, developing a more comprehensive understanding of *Mh* ecology is critical. *Mh* has been characterized based upon serotype, and more recently genotype, with serotypes 1 and 6 traditionally considered to be more associated with disease^13,14^. More recently, a WGS-based study of over 1,000 isolates revealed two dominant genotypes of *Mh* with distinct associations to disease status^2^, highlighting the potential importance of genomic variants in the ecology of this agent. These important findings regarding ecology and pathogenicity of *Mh* have been completely reliant on the ability to culture the organism to enable further characterization, which inherently limits the ability to investigate targets within the entire microbial population. To date, culture-independent investigations of *Mh* have largely been limited to 16S rRNA sequencing, with relatively few studies using shotgun metagenomics with limited sequencing depth ^15–18^. As a result, these data have been unable to resolve variant-level diversity within *Mh* communities. Together, these insights underscore the need for a framework capable of producing sufficient sequencing information to enable characterization of variant-level diversity directly from metagenomic data.

Here, we have developed and validated a comprehensive molecular and computational workflow to characterize variant-level diversity within microbial populations. Notably, we used TE metagenomic sequencing to greatly increase the on-target shotgun sequencing efficiency. We validated this workflow by assessing variant-level *Mh* populations in the respiratory tract of cattle with differing loads of *Mannheimia* spp. and respiratory disease status. Importantly, we developed a variant-level classification system for *Mh* based on all publicly available genomes in NCBI’s GenBank and RefSeq database at the time of writing (n =2,435; January 2025). To address the ambiguity associated with the term “strain,” we introduce genomically clustered sequence variants (GSVs) as a useful sub-species discriminatory and biologically relevant classification unit. These GSVs represent distinct clusters created from the analysis of genome-wide sequence similarity. Our accompanying bioinformatic workflow, VARIANT++, developed to enable this analysis, is easily implemented and highly adaptable for analysis of TE metagenomic data targeting any microbe of interest.

## Results

### Study overview

To analyze variant-level dynamics of a target species within a diverse microbial population, we developed an integrated molecular laboratory and bioinformatic workflow. We designed a set of biotinylated RNA baits based on all publicly available reference genomes and used them to perform target-enriched (TE) shotgun sequencing. We developed a novel annotation scheme to enhance variant-level classification by developing an approach that leverages analysis of genomic similarity in creating a reference database of GSVs, which is used in conjunction with a custom bioinformatic pipeline, VARIANT++ (https://github.com/Microbial-Ecology-Group/VARIANTplusplus). This bioinformatic processing includes QC trimming, read merging, host DNA removal, and read filtering using a k-mer classifier to assign species-level classification to the reads. These species-level reads are then processed for GSV classification using Themisto ^19^ for pseudoalignment to reference genomes and mSWEEP ^20^ to statistically infer the relative abundance of counts between closely related GSV groups.

To demonstrate a practical application of this workflow, we targeted the important cattle pathogen, *Mannheimia haemolytica*, using samples collected from beef cattle. To optimize target-enrichment, we investigated different sequencing approaches using a set of “Optimization” samples to inform further the sample preparation methodology. We then applied the optimized method in the characterization of another set of “Investigation” samples, consisting of nasal swabs collected from feedlot cattle with differing relative abundances of *Mannheimia*, based on 16S rRNA gene amplicon sequencing data.

### Bait design for target enrichment

#### Bait design

Using 70 *Mh* reference genomes, previously described tools were used to create a parsimonious set of cRNA DNA template baits that align across the *Mh* genome. Our bait design consisted of 48,394 sequences for the target enrichment of *Mh* (Supplementary File 1). To evaluate the ability of the bait set to detect other *Mh* sequences, these baits were aligned against a randomly selected *Mh* genome not used in the bait design (M42548, RefSeq Accession: NC_021082.1). An average of 2× coverage across the genome was calculated (range: 1-16× per base) with a percent identity of 92% across the entire genome (2,731,870 bp). Additionally, we aligned the baits to a representative genome from each of the GSV groups described below, which identified an average 97.7% pairwise identity (range: 95.8% – 98.9%) and an average 1.9× genome coverage depth (range: 1.7× – 2.1×) for all GSV groups (Supplementary Table 1). Together, this suggests adequate coverage across *Mh* genomic regions with sufficient allowance for mismatches, suggesting that they operate within the manufacturer’s guidance that baits can perform acceptably with as much as 20% mismatch in the target regions. Finally, balanced bait boosting was applied using Agilent’s eArray software, to improve pull-down performance by increasing the representation of both orphan (“no immediate neighbors in the genome within 1 bp on either side of the bait”) and GC-rich baits. The resulting *Mh* bait set (n=114,375) was used for hybridization and sequence capture during library preparation (Agilent design ID S3319212).

#### Optimizing target enrichment performance

To determine the optimal protocol for enriching *Mh* within metagenomic sequencing libraries, five samples with high relative abundances of *Mannheimia* genus determined using 16S rRNA gene sequencing were purposively selected (Optimization Samples). For each sample, isolated DNA was used to generate ten technical replicates that were used to investigate five different modifications of the target-enrichment (TE) procedures in parallel. One method performed the hybridization and target enrichment process using the manufacturer’s recommendation for probe quantity (FULL). Because we hypothesized that the molarity of baits in relation to target sequences can affect capture efficiency, two modified enrichment processes used 50% and 25% of the recommended baits amount for the hybridization process (HALF and QUARTER, respectively). Our prior experience suggests that the abundance of target sequences can influence the pull-down yield when off-target DNA is very abundant, the fourth and fifth methods performed the hybridization and enrichment process twice in succession, each step using 25% or 50% of the recommended bait quantity, before completing the library preparation (DOUBLE QUARTER and DOUBLE HALF, respectively). Thus, the five protocols can be grouped into single-capture methods using varying bait amounts (FULL, HALF, QUARTER) and double-capture methods using reduced bait in two successive rounds (DOUBLE HALF, DOUBLE QUARTER). For comparison, the same five metagenomic samples also underwent deep shotgun sequencing using traditional methods.

Sequencing of the optimization samples generated an average of 71,846,997 paired end reads across the 39 TE sequencing libraries (range: 1,170,835 – 153,793,168) (Supplementary File 2) and an average of 131,233,940 paired-end reads across the 5 shotgun sequencing libraries (range: 120,365,961 – 141,073,067)(Supplementary File 3). After quality-filtering and the merging of overlapping reads, replicates sequenced with traditional shotgun methods did not differ from each other in read depth (Fig. 1a; pairwise Wilcoxon rank-sum, n = 5-9, p < 0.05), but shotgun sequencing produced significantly more total reads than the 5 TE protocols (Fig. 1a; pairwise Wilcoxon rank-sum, n = 5-9 per protocol, p < 0.05). This was likely because shotgun and TE library prep and sequencing were performed separately and with differences in target sequencing depth. We observed higher rates of duplicate reads in TE libraries (mean: 84.92%, range: 54.4% - 95.2%) compared to shotgun (mean: 21.72%, range: 19.58% - 22.76%). After deduplication, a similar trend was observed where shotgun libraries retained significantly more reads than all 5 TE libraries (Fig. 1b; pairwise Wilcoxon rank-sum, n = 5-9, p < 0.05). Within TE libraries, FULL retained significantly more reads after deduplication than the other four TE protocols (Fig. 1b; pairwise Wilcoxon rank-sum, n = 5-9 per protocol, p < 0.05). However, the removal of host-aligned reads resulted in a dramatic drop in read counts for traditional shotgun libraries, which yielded significantly fewer non-host reads than DOUBLE HALF TE libraries (Fig. 1c; pairwise Wilcoxon rank-sum, n = 5-9, p < 0.05). While the median numbers of non-host reads were much greater for both double-capture methods (DOUBLE QUARTER and DOUBLE HALF), the greater variability in yields from QUARTER and DOUBLE QUARTER libraries suggests that lower bait concentration can affect uniformity of capture yields, and the variability affected the ability to demonstrate statistical differences in median counts (Figure 1a-c).

**Figure 1.**
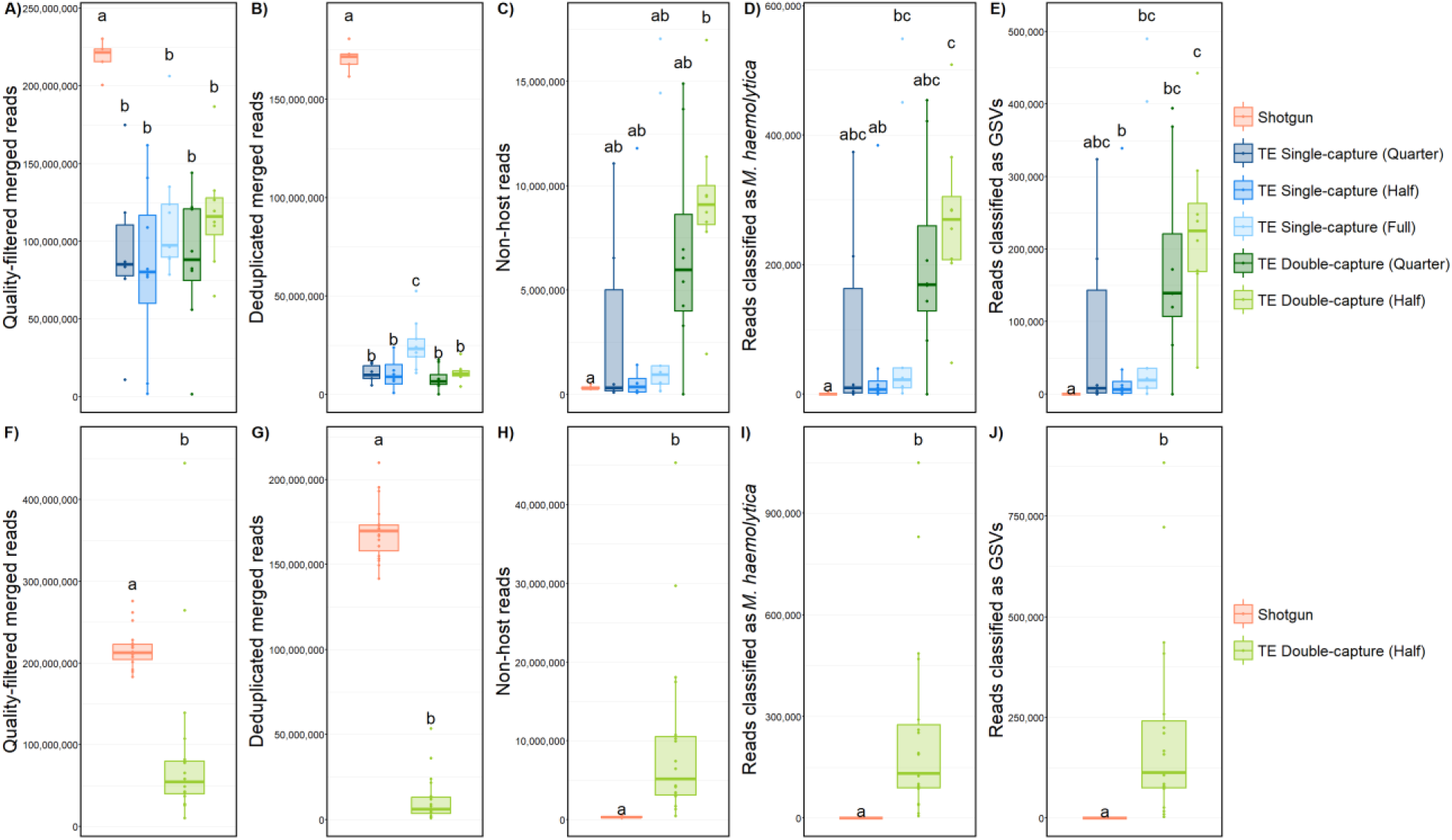
Performance of target-enrichment (TE) capture protocols versus shotgun sequencing across two sample sets. Panels A–E show the Optimization samples; panels F–J show the Investigative samples. Columns share the same response variable: (A,F) quality-filtered & merged reads; (B,G) deduplicated merged reads; (C,H) non-host reads after host depletion; (D,I) reads classified as *M. haemolytica*; (E,J) reads classified as genomic sequence variants (GSVs). Boxplots summarize distributions across libraries (center line = median; box = IQR; whiskers = 1.5×IQR; points = individual libraries). Colors denote library/capture protocols (see legend): Shotgun; TE single-capture (Quarter, Half, Full); TE double-capture (Quarter, Half). Different letters above boxes indicate groups that differ significantly (pairwise Wilcoxon rank-sum test, α = 0.05); groups sharing a letter are not significantly different.

Critically, extremely large differences in median on-target counts were observed when evaluating the number of reads classified as *Mh*. The DOUBLE HALF protocol produced, by median, more than four-fold greater *Mh*-aligned reads than FULL, HALF, or traditional shotgun protocols (Fig. 1d; pairwise Wilcoxon rank-sum, n = 5-9 per protocol, p < 0.05). The median number of *Mh*-aligned reads was also greater for FULL and HALF single-capture protocols when compared to traditional shotgun sequencing, though this was only statistically significant for the FULL protocol. While the DOUBLE QUARTER protocol also yielded a greater median number of *Mh*-aligned reads than the single capture or shotgun protocols, as noted previously, variability in total counts from QUARTER and DOUBLE QUARTER protocols was substantial, affecting statistical comparisons. These samples were also analyzed with our workflow described below, to further assign reads classified as *Mh* to the GSV level. While only 1 out of 5 shotgun samples had non-zero GSV counts, similar patterns were observed with the DOUBLE QUARTER protocol yielding a higher median number of reads classified as GSVs compared to single capture, with a high variability in the number of reads from QUARTER and DOUBLE QUARTER protocols (Fig. 1e; pairwise Wilcoxon rank-sum, n = 5-9 per protocol, p < 0.05).

### Optimization of the classification workflow

#### Taxonomic classification of Mannheimia haemolytica

Several preprocessing steps influence the performance of downstream classification accuracy, including read-quality trimming, depletion of host DNA and taxonomic assignment. Therefore, our workflow started with QC trimming using Trimmomatic^21^. We then merged reads with FLASH^22^ and deduplicated them with seqKit^23^ to increase average read length and account for the potential of PCR bias, respectively. QC-trimmed, merged, and deduplicated reads were then processed for host removal with bwa^24^ and samtools^25^.

We tested various classification tools and databases for taxonomic assignment, electing to use Kraken2^26^ paired with a comprehensive database, the core NT database (https://benlangmead.github.io/aws-indexes/k2), which contains all NCBI GenBank reference genomes. This, paired with increasing the “--confidence" flag, reduces false positive classification because it will classify reads based on least common ancestor (LCA), and the large database allows for comparison to more genomes. Alternatively, including all GenBank *Mh* genomes increases the number of classified reads because we can identify unique regions in as many *Mh* genomes as possible. We also evaluated using a smaller database and adding all available *Mh* genomes, which increases the number of *Mh* classified reads by highlighting unique regions; however, including genomes from other, “off-target” taxa improve overall classification confidence by allowing the algorithm to identify similarities across related species. However, classification past the species level is challenging with this tool; therefore, we developed and optimized a database that could be used to further characterize reads classified as *Mh* using LCA in relation to objectively (data-driven) classified genomic variants based on average nucleotide identity (ANI). To achieve this, we grouped published *Mh* genomes into genomic sequence variant (GSVs) clusters based on whole-genome sequence similarity. Only reads classified as *Mh* at the species level were extracted and used for downstream GSV analysis.

#### Using simulated samples to optimize GSV classification

To create our custom GSV database, all *Mh* genomes from GenBank and RefSeq as of January 2025 (n=2,435) were included. Pairwise whole-genome similarity, using an all-versus-all approach, was measured using ANI with FastANI^27^. Then, because the optimal number of variant-level GSV groups for best distinguishing between closely related genomes was unknown, we ran benchmarking tests consisting of simulated samples generated by subsampling reference genomes and processing them through our classification workflow with an increasing number of unique GSV groups (2-30). Additionally, we tested differences in sequencing depth, Kraken2 confidence scores, GSV richness in a sample, and evaluated false-positive classification proportions using off-target genomes from other species in the Pasteurellaceae family.

The results from analyzing simulated *Mh* reads with GSV annotation schemes of increasing cluster number were plotted and evaluated for the best precision and recall (Supplementary Figs. 1-3). Based on the simulation data, we determined that false-positive classifications were reduced when filtering to only include reads classified as *Mh* at the species level using LCA, and that the annotation scheme with 8 GSVs performed optimally (recall = 91.3%, precision = 96%), as there was a statistically significant drop in performance as the number of GSV groups was increased (Supplementary Fig. 3). The final reference database annotation scheme was primarily dominated by two large groups of 1,626 and 691 genomes, with the remaining 118 reference genomes distributed across the 6 other GSV groups. The average ANI within genomes in the same GSV groups were above 99.28%, except for GSV 6, which had a lower average of 96.7% within-group ANI (range: 96.7% – 99.9%)(Supplementary Table 2). A representative genome was identified from each GSV group using dRep ^28^, and pairwise distances were summarized between all GSV groups (Supplementary Table 3). GSV 6 was the most divergent group, with all pairwise ANI values below 96%; by contrast, all other GSV groups were at least 98.14% similar to each other.

Considering the previous genotypic classification of reference isolates^5^ that is widely used in studies of *Mh*, the GSV cluster containing genomes previously classified as Genotype 1 was arbitrarily designated as GSV 1, and the GSV cluster containing genomes previously classified as Genotype 2 was designated GSV 2 (Fig. 2). Interestingly, one genome previously classified as being “between genotype 1 and genotype 2”^29^ was clustered separately in our GSV analysis, and was designated GSV 4. The similar separation of previously classified genomes while also identifying additional genomic variants emphasizes the relevance our new classification while providing additional granularity. This final database of 2,435 *Mh* genomes and their 8 GSV annotations was used for downstream analysis.

**Figure 2.**
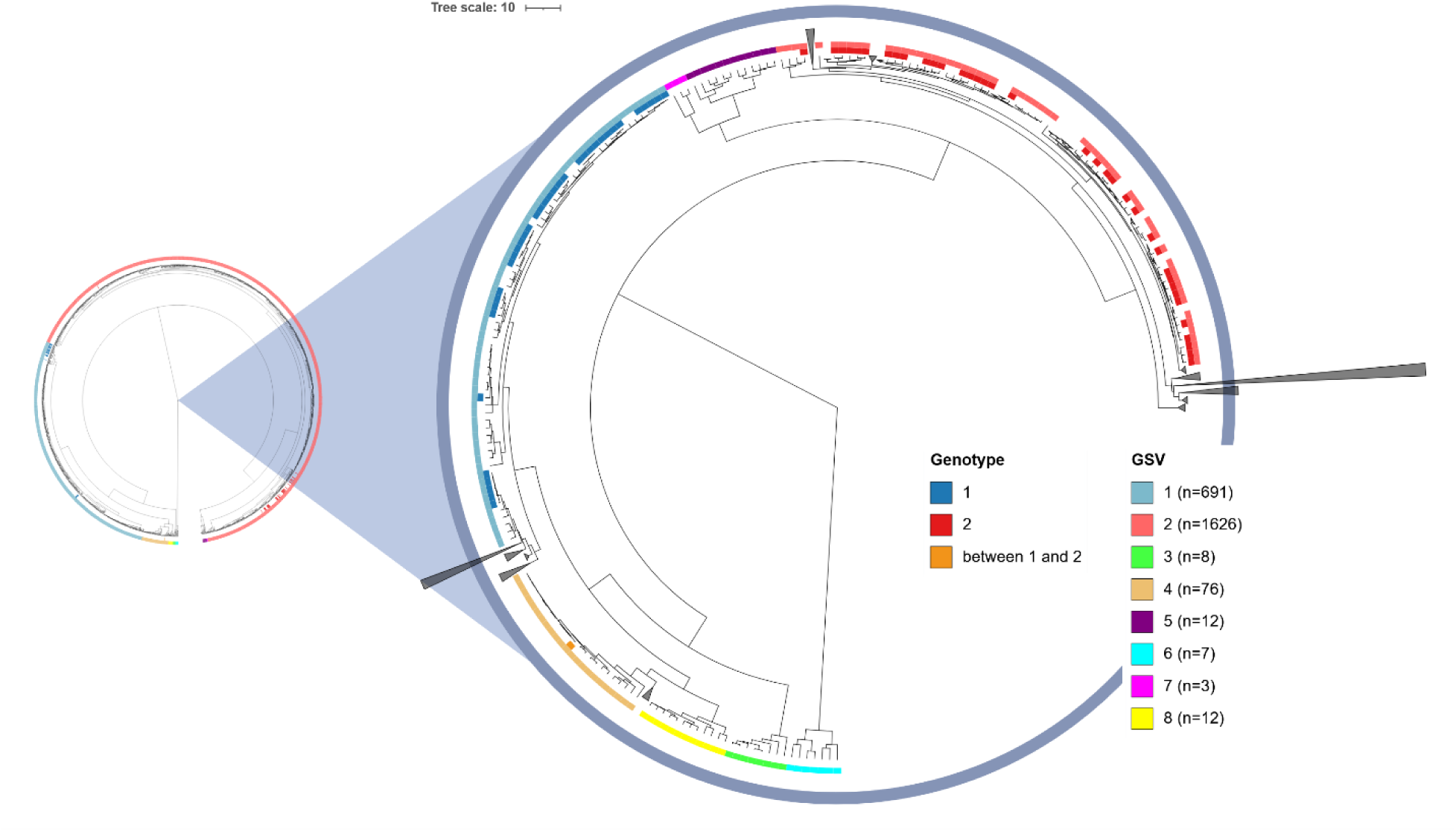
ANI distance tree of 2,435 *Mannheimia haemolytica* genomes with genotype and GSV annotations. Left: full circular tree. Right: magnified region showing tip annotations. Branch lengths are proportional to ANI-derived distances (scale bar = 10). Two concentric tip tracks are shown: Genotype (inner track; blue = 1, red = 2, orange = “between 1 and 2”) and GSV group (outer track; GSV 1–8 with sample counts shown in the legend). In the zoomed panel, grey triangles indicate collapsed monophyletic clades that contained only a single GSV and no genotype-labeled isolates. Overall, genotype labels broadly align with GSV groupings, large genotype 1 segments coincide with GSV1-enriched clades and genotype 2 segments with GSV2-enriched clades, while isolates labeled “between 1 and 2” occur near boundaries of these clades, consistent with intermediate ANI relationships. Trees were exported from iTOL and the final figure was created in BioRender. Doster, E. (2025) https://BioRender.com/iblmpn8

Overall, benchmark analysis identified a low proportion of false-positive classification in all mock samples created from off-target Pasteurellaceae genomes. Using the default Kraken2 settings led to an average of 303 reads incorrectly classified as *Mh* out of an average 907,978 input reads per sample, and rates decreased as the Kraken2 confidence score value was increased. Given the varying false-positive rates across GSV groups, and the superior performance of the 8-GSV annotation scheme in our tests, we centered the analysis on the 8-GSV annotations. To conservatively account for false-positive classification, we calculated the 99^th^ quantile for the proportions of false-positive counts for each GSV as classified with a Kraken2 confidence score of 0 (GSV 1: 0.0000653, GSV 2:0.00138, GSV 3:0.000743, GSV 4:0.000739, GSV 5:0.000427, GSV 6:0.00104, GSV 7:0.000283, GSV 8:0.000765). Thus, to balance GSV classification performance and false positive proportions, we elected to use Kraken2 with a confidence score of 0.1 to extract reads classified as *Mh* and, following GSV classification with Themisto and mSWEEP, apply a count filter based on the expected false positive rate of each GSV as estimated from the benchmark analysis. This approach prioritizes minimizing false positives, even if it means some low-abundance GSVs may be missed, though other users can adjust the confidence threshold and filtering strategy according to their own analysis goals.

#### Development of the VARIANT++ bioinformatic pipeline

We developed the VARIANT++ bioinformatic pipeline using the Nextflow DSL2 programming language^30^ to facilitate this optimized variant-level classification of *Mh* employing our novel GSV database annotation scheme (https://github.com/Microbial-Ecology-Group/VARIANTplusplus). This nextflow-based pipeline processes short-read metagenomic sequences for QC trimming, read merging, read deduplication, host DNA removal, and taxonomic classification with Kraken2. Reads classified as *Mh* are then extracted and used for downstream GSV analysis. Extracted reads are classified using Themisto and our custom *Mh* database, followed by GSV classification using mSWEEP. A count filter is then applied, using the calculated false positive rate for each GSV, calculated by multiplying the false positive rate by the number of non-host reads and subtracting the product from the counts for that GSV. Instructions on the GitHub page can be followed to re-create this workflow on any species of interest (https://github.com/Microbial-Ecology-Group/VARIANTplusplus/blob/master/docs/VARIANT%2B%2B_step_by_step.md).

### Application to the Investigation Samples

#### Investigation samples

To further evaluate these novel molecular and bioinformatic workflows, we selected a subset of “Investigation” samples from those collected as part of a previously published study of feedlot cattle^16^ (Supplementary File 4). These samples (n=120) were stratified into five categories based on their relative abundance (RA) of *Mannheimia* genus-level reads that was previously estimated using 16S rRNA gene sequencing results: 1) no *Mannheimia* genus-level reads detected (Zero); 2) >0% and ≤1% RA (Low); 3) >1% and ≤10% RA (Medium), 4) >10% and ≤30% RA (High), and 5) >30% RA (Highest). We then used stratified random sampling to select the samples to be evaluated using TE shotgun sequencing as an example investigation of *Mh* GSVs that can be found in feedlot cattle (Investigation Sample Set; n = 4 for each abundance stratum, total n = 20).

The DOUBLE HALF protocol was selected as being optimal for the analysis of the Investigation sample set. Using TE, shotgun libraries were successfully prepared using this protocol for 19 swab samples; one sample from the “Low” *Mh* stratum failed during pre-capture PCR in the initial attempt and was not attempted again. Sequencing of these 19 TE shotgun libraries produced an average of 58,732,198 paired-end reads per sample (range: 7,386,424 – 295,111,085; Fig. 1), with no identifiable differences in sequencing depth among the *Mannheimia* relative abundance categories (pairwise Wilcoxon rank-sum, n = 3-4 per category, p > 0.05) in the number of raw sequencing reads, non-host reads, reads classified as *Mh*, or the number of reads classified as GSVs (pairwise Wilcoxon rank-sum, n = 3-4 per category, p > 0.05). Classification using the VARIANT++ pipeline and our GSV database identified reads attributed to 6 unique GSVs among the 19 samples, with an average of 250,134 (2.51% of non-host) reads per sample classified to *Mh* GSVs (min: 6,594, max: 1,049,743; Fig 1; Supplementary File 2-4). The *Mh* population in these samples was largely dominated by GSV 2, which was present at an average relative abundance of 97.4% across all samples (Fig. 3 – GSV heatmap). The reads classified to other GSVs were sparse and in low abundance, with only GSV 4 and 6 not present in these samples (Fig. 4– prevalence plot).

**Figure 3:**
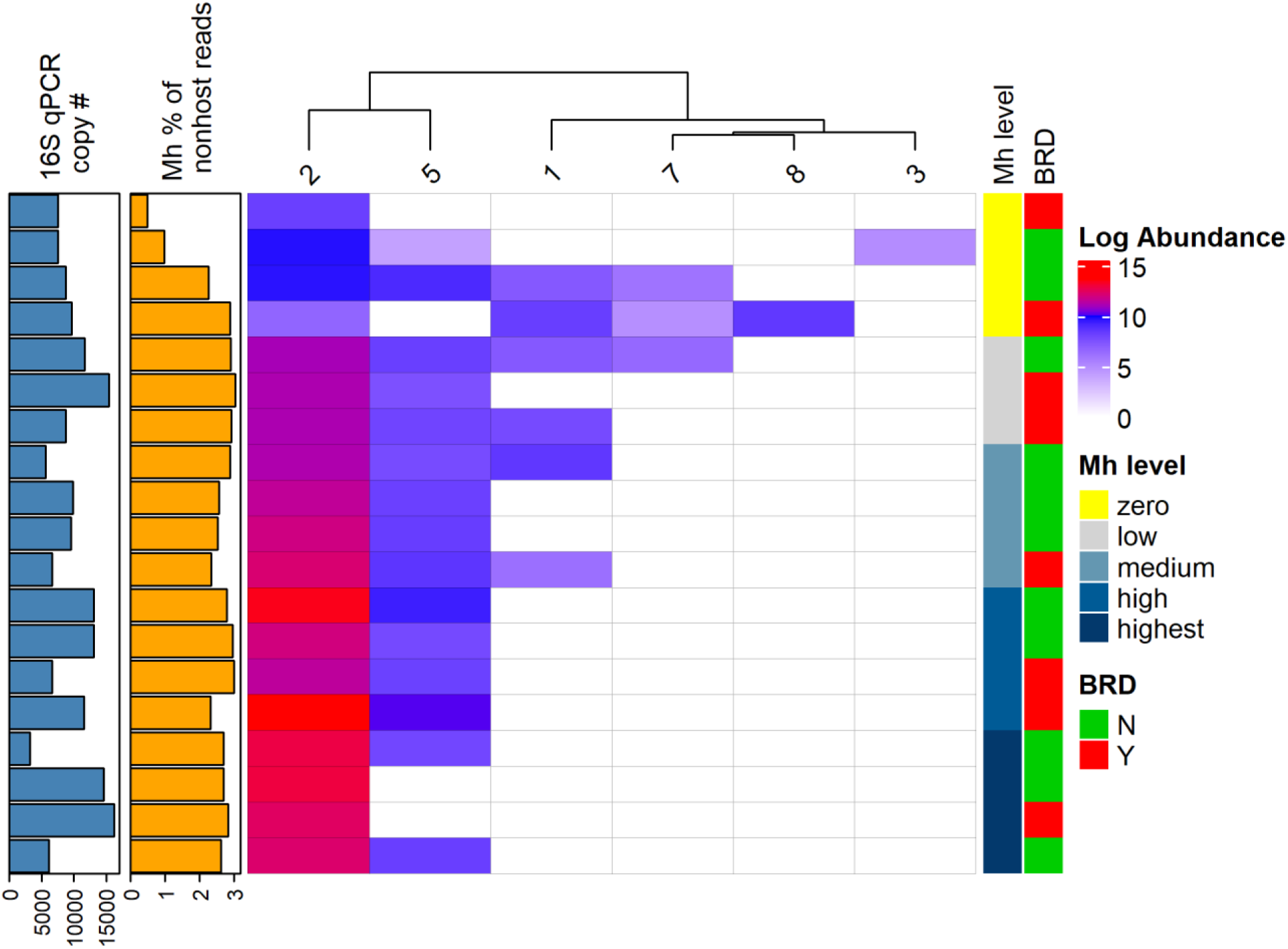
GSV abundance heatmap with per-sample metadata. Rows are samples; columns are M. haemolytica genomic sequence variants (GSVs; column dendrogram shows hierarchical clustering of GSVs by abundance profile). Heatmap cells show log-transformed counts for each GSV. On the left, two horizontal barplots summarize per-sample covariates (same row order as the heatmap): “16S qPCR copy #” and “Mh % of nonhost reads” (percent of nonhost reads classified as *M. haemolytica*). On the right, row annotations indicate *Mh* level strata, and BRD status.

**Figure 4:**
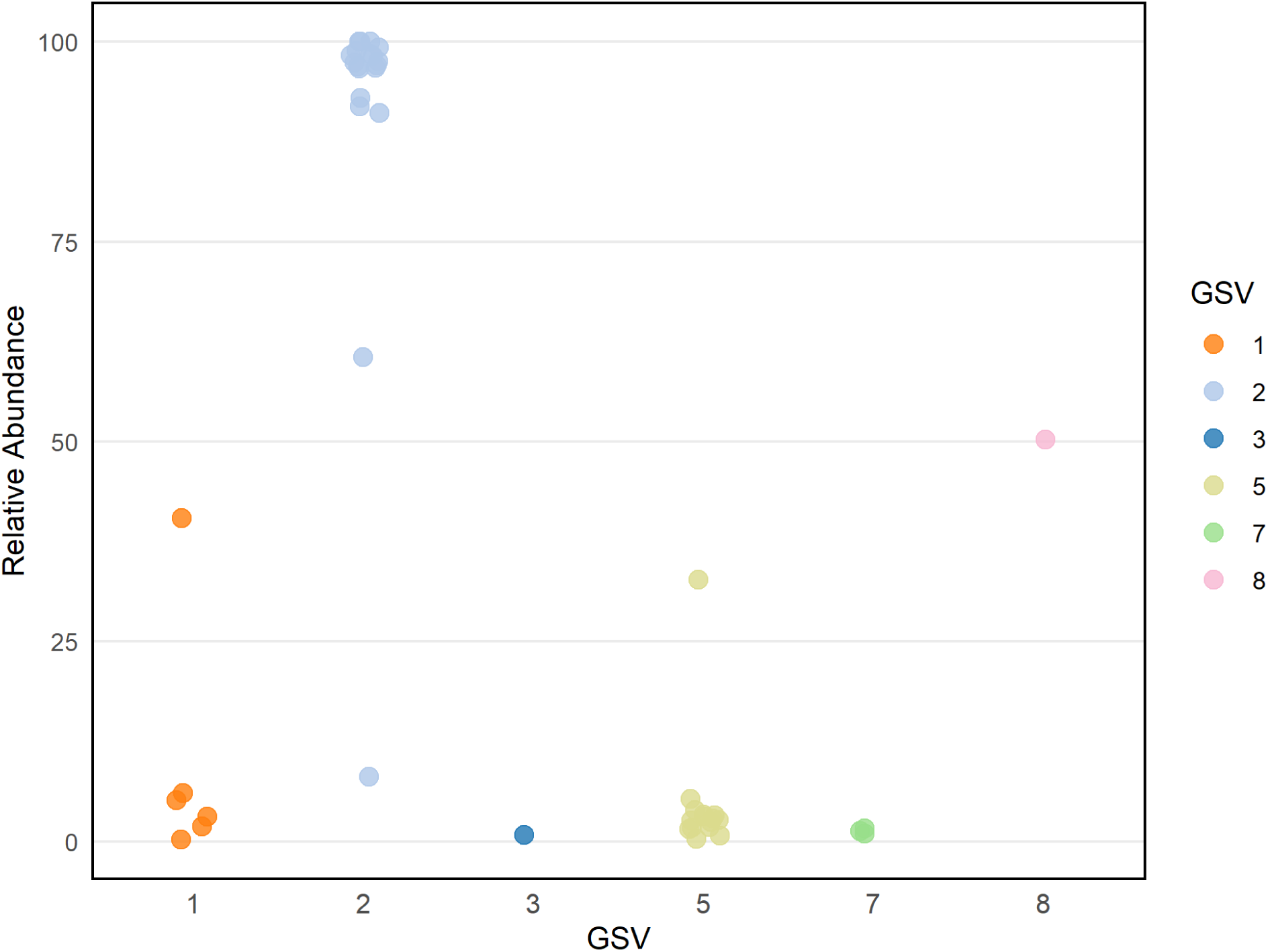
Relative abundance of *M. haemolytica* genomic sequence variants (GSVs) across samples. Each point represents the relative abundance of a GSV in one sample; the x-axis is the GSV (1–8) and the y-axis is the within-sample relative abundance (%). Points are colored by GSV identity and horizontally jittered to reduce overplotting. Of the eight predefined GSVs, only GSV-4 and GSV-6 were not detected in this dataset.

These same 19 samples from the Investigation Sample Set were also sequenced using traditional shotgun sequencing and analyzed using VARIANT++. Libraries generated from TE sequencing had significantly fewer QC-trimmed reads and merged-deduplicated reads compared to libraries generated from shotgun sequencing (Fig. 1f-g; pairwise Wilcoxon rank-sum, n = 19 per protocol, p < 0.05). TE libraries had higher read duplicate rates (mean: 87.36%, range: 69.65% - 93.49%) compared to shotgun (mean: 21.89%, range: 19.41% - 25.39%), however, TE sequencing resulted in significantly higher counts of non-host reads, reads classified as *Mh*, and reads classified as GSVs (Fig. 1i-j; pairwise Wilcoxon rank-sum, n = 19 per protocol, p < 0.05). While all 19 samples had non-zero counts for *Mh* classification at the species level, the average count was 656 (range: 1 - 4297) across all shotgun samples, which limited the ability to perform GSV analysis. As a result, only 5 out of 19 shotgun samples were positive for GSVs after accounting for expected false-positive read proportions, and all were attributed to GSV 2. Interestingly, only samples in the “HIGH” or “HIGHEST” *Mh* strata generated shotgun metagenomic reads that could be classified for GSV counts.

In the TE samples, GSV composition was found to differ significantly by Mh strata (PERMANOVA R² = 0.54, P = 0.0001). Before accounting for multiple comparisons, the GSV composition of samples in the ZERO *Mh* strata was significantly different than those in the LOW (PERMANOVA R^2^=0.44, P = 0.03), MEDIUM (R^2^=0.43, P = 0.03), HIGH (R^2^=0.34, P = 0.03), and HIGHEST (R^2^=0.5, P = 0.03) strata. After applying the Benjamini-Hochberg correction, these differences were no longer statistically significant (P < 0.05). Although the analysis lacked sufficient power to detect significant differences, visual inspection suggests a trend indicating that samples in the ZERO stratum differed from those in higher strata (Supplementary Fig. 4 – NMDS). This trend is primarily driven by the identification of multiple GSVs in the ZERO *Mh* strata, including those other than the dominant GSV 2 and an overall decrease in GSV counts (Fig. 3 – GSV heatmap). However, there were no significant differences in beta diversity composition of GSVs when stratified by BRD status.

## Discussion

Shotgun metagenomic and marker-based sequencing are widely recognized for providing insight into microbial communities without the selection bias inherent to culture^31,31,32^. Our work demonstrates the substantial gains in analytical sensitivity and specificity achievable with target-enriched (TE) shotgun sequencing, enabling in-depth characterization of microbial populations at the sub-species level^3,33,34^. This approach allows for more comprehensive interrogation of microbial taxa in complex metagenomic samples than standard shotgun or 16S amplicon sequencing. Importantly, the method is adaptable to any taxa of interest, taking advantage of the continually expanding catalog of reference sequences.

Using *Mh* as a case example, we have shown that this approach will generate detailed information about genomic variants that can be present even within individual samples. This novel approach therefore provides insights regarding microbial ecology of important pathogens at a resolution and scale previously unattainable with other approaches, fostering a deeper understanding of disease pathogenesis. Specific to respiratory disease in cattle, which is the most costly disease of beef cattle and the most common illness treated with antimicrobial drugs, the number of classified GSVs for *Mh* detected across all samples suggests that there is a greater degree of genomic variability than was previously recognized in this important pathogen of cattle^5,35^.

Importantly, our molecular and bioinformatic workflows are likely adaptable to investigate any other target within metagenomic microbial communities from any ecosystem. It is also suitable for TE metatranscriptomic sequencing, and with greater modifications, the approach could be utilized with long-read sequencing. Our bioinformatic workflow can also be applied to traditional shotgun sequencing of single genomes or metagenomic communities to classify at the sub-species level using a data-driven taxonomy based on whole-genome sequence similarity (i.e. determined from ANI analysis). However, the target-enriched molecular workflow described here provides exponentially greater depth of interrogation of the target, resulting in greatly increased interrogation power with similarly enhanced resolution. It is notable that the method for classification of genomic variants described here circumvents issues associated with arbitrary sub-species annotations (e.g., what constitutes a strain?) that can often be found in the literature and carries over to public genomic repositories. Annotations uploaded with sequence information can be inconsistent at the strain level and sometimes appear arbitrary. Further, they can change over time and vary depending on the original purpose of sequencing each genome.

Our approach for clustering genomes using ANI to objectively characterize genomic similarity addresses this issue, providing an objective and quantifiable method for annotating and analyzing variant-level categories of any microbial taxa. Importantly, the resulting database serves as an external reference that is not subject to the internal validity constraints of databases created with operational taxonomic units (OTUs): new genomes can be added, and unless they are highly divergent, they will typically cluster within the existing GSV framework. Our benchmarking with *in silico* samples was important for informing numerous analytical options, creating a flexible workflow that can be optimized based on user needs. For example, while we elected to prioritize the reduction of false-positive classifications, the workflow could be easily tailored to fit other research priorities, such as maximizing sensitivity. Another factor influencing classification performance is k-mer size; while we did not test optimization of this parameter for Kraken2 or Themisto pseudoalignment, the developers of these tools performed extensive benchmarking to recommend default k-mer sizes^19,26^. Each tool performs its respective task effectively, but we believe that our multi-step combination of tools and filtering strategies is key to optimizing classification in target-enriched data. Although the full phylogenetic scope of variant diversity (i.e., the number of biologically meaningful GSV groups) remains unresolved, our workflow uses established tools to provide robust evidence for subspecies-level classification in this study.

Exploring various protocols for preparing TE shotgun libraries was also important for creating optimized molecular methods that are essential to this integrated workflow. Despite having double the sequencing depth using traditional shotgun methods (Fig 1), the logarithmic improvement in on-target sequencing using TE methods was critical to the success of this approach. With these respiratory swab samples, TE sequencing overcame both an abundance of background host DNA as well as highly diverse metagenomic communities. In these circumstances, the DOUBLE HALF protocol yielded more than four times the on-target sequencing depth at the most refined classification resolution when compared to single capture TE protocols. It appears that using the QUARTER and DOUBLE QUARTER protocols amplified the variability in on-target sequencing yields. We have successfully used the HALF protocol when performing TE shotgun sequencing of rare targets within highly diverse microbial communities, such as resistance genes in fecal samples ^10,11^, and our results suggest that the double enrichment protocols may not be as important in the absence of a highly abundant background such as the host DNA found in the nasal swabs used in this investigation. However, the significant portion of duplicated reads generated may suggest that refining PCR conditions may further optimize these protocols. The authors have noted libraries that pass QC prior to hybridization and capture that go on to fail after the post-capture amplification in other projects. Sometimes, re-running the hybridization and capture steps with more pre-capture library input results in a successful library prep. In other cases, the final library fails despite different attempts. Because our goal was to optimize the TE process in this project, we decided to note the library prep fails and not include them in the sequencing pool.

The development of *Mh*-TE, our GSV annotated *Mh* database, and the VARIANT++ bioinformatic pipeline enhanced the precision and efficiency of variant-level characterization from metagenomic communities without the need for culture or whole genome sequencing. This allows us to better characterize the microbial ecology of *Mh* populations while avoiding selective biases that are inherently associated with culture. While culture-based methods and whole-genome sequencing can provide a wealth of detailed information on genomic variability of microbial taxa, this novel approach paves the way for more comprehensive studies of bacterial populations within highly diverse metagenomic communities. Ultimately, this new perspective of the ecology of varied bacterial populations within an individual taxon may provide important new insights regarding the epidemiology, pathogenesis, and control of important pathogens such as *Mh*.

## Methods

### Bait design

A novel bait set was designed for variant-level *Mannheimia haemolytica* (*Mh*) differentiation using 70 reference *Mh* genomes (NCBI Accessions: CP017484-CP017552 and CP006957). First, a primary 120-mer bait set was generated using the program CATCH^36^ and the parameters “--cluster-and-design-separately 0.5 -e 50 -pl 120 -ps 120 -m 20”. This initial design yielded 51,612 probes. The bait set was then refined using Kraken2 to verify *Mh* specificity, which resulted in 47,011 probes specific to *Mh*. To enrich integrative conjugative elements (ICE) associated with *Mh*, we aligned three specific ICE sequences (NCBI accessions: CP004752, CP005383, and CP017551) from *Mh* whole-genome sequences to NCBI’s “nt” database using ‘blastn’ within BLAST+ version 2.10.0^37^ to identify additional ICE sequences. From this, we extracted 399 sequence variants with over 80% identity to reference ICE elements, leading to the creation of an extra 2,131 probes. Further, to enhance the probe set’s utility in discriminating *Mh* genotypes, we selected 13 informative loci within the *Mh* genome and identified reference sequences to target these regions (Supplementary File 5). We then cross-referenced these loci against the unfiltered bait set using Geneious software^38^ (2021.2 Dotmatics), incorporating an additional 1,361 probes that aligned with these regions.

Bait sequences from these three sources were combined, and CD-hit^39^ was used to remove redundant sequences by clustering at 98% sequence identity, resulting in 48,394 baits. To evaluate the sensitivity for the baits to recognize sequences from *Mh* genomes not in the bait design, the final bait design (n=48,394) was then aligned using the Geneious mapper with default settings to a randomly selected *Mh* genome (M42548, RefSeq Accession: GCF_000376645.1). To assess the bait coverage across different GSV groups in a given annotation set, we used dRep and the “dereplicate” function^40^ on each group of GSV genomes to identify a representative genome. We then aligned the baits to each representative genome to summarize the pairwise identity and mean coverage across the entire reference sequence. Representative genomes that did not have a completely assembled sequence, were first assembled using the Geneious assembler with default settings, and coverage was summarized across all resulting contigs.

Bait boosting added 65,981 baits for a final *Mh* bait design set with a total of 114,375 baits. Target-enriched sequencing libraries were prepared using the SureSelect XT HS2 Reagent Kit for Illumina Paired-End Multiplexed Sequencing (Agilent Technologies), with modifications (Supplementary text in GitHub: https://github.com/Microbial-Ecology-Group/Manuscript-Mh_TE_validation/blob/main/Methods/VERO%20Target%20Enriched%20Single%20%26%20Double%20Capture.docx).

### Study samples

#### Population

Samples used in this study were collected to investigate methods for the detection and characterization of *Mh* and the upper respiratory microbiome and resistome of cattle, which has been partially described previously^16^. Briefly, beef-type steers and bulls (n=120) were purchased from a commercial livestock auction and shipped to the West Texas A&M University Research Feedlot. Cattle weighed an average of 261.2 kg (SD = 12.2 kg) at arrival and were fed for an average of 235 days (range 213-255 days) prior to harvest. Upon arrival, all animals were vaccinated against respiratory and clostridial pathogens, treated with anthelmintics, and treated with tildipirosin (4 mg/kg subcutaneously, Zuprevo, Intervet Inc., Summit, NJ), a long-acting macrolide, as methaphylaxis for BRD. Cattle were randomly assigned to pens (n=8 per pen) where they were housed and fed until harvest. Cattle were monitored daily through the feeding period by trained feedlot personnel, and animals with moderate to severe clinical signs of respiratory disease and a fever (rectal of temperature ≥40°C) were classified as having BRD and treated with antimicrobial drugs under the supervision of a veterinarian. Overall, 36.4% of the cattle were diagnosed with BRD and treated during the feeding period, with a median of 10 days until first treatment. Protocols for animal use related to this research were reviewed and approved by the West Texas A&M University Institutional Animal Care and Use Committee (Protocol# 2020.04.003).

#### Samples

Samples used in this research were collected from study cattle 14 days after arrival. Briefly, a16-inch (40.6 cm) proctology swab with an oversized rayon fiber tip (816-100, Puritan, Guilford, ME) were passed through the nares to the caudal limit of the nasopharynx at the level of the palatopharyngeal arch, as previously described^16^. Swabs were collected aseptically, and tips were cut into sterile 5ml tubes with 100% ethanol to stabilize microbial populations, placed on ice, and transported within 2 hours of collection to the laboratory where they were frozen at – 80°C until further processed. From these samples, two sets were purposively selected as “Optimization” and “Investigation” samples to test sample preparation methods to improve on-target sequencing and to test our entire workflow, respectively.

#### DNA isolation

DNA was isolated from all samples using a commercial extraction kit (QIAamp Power Fecal Pro DNA, Qiagen, Hilden, Germany) according to the manufacturer’s instructions. Following isolation, DNA was quantified (ng/µL) using fluorometry (Qubit Flex, Thermo Fisher Scientific).

#### 16S rRNA gene sequencing and analysis

All samples were evaluated using 16SrRNA gene sequencing, as previously described^16^. Briefly, the V3-V4 region of the 16S rRNA gene was amplified using the 341F (5’ – CCTACGGGNGGCWGCAG – 3’) and 805R (5’ – GACTACHVGGGTATCTAATCC – 3’) primer pair (Integrated DNA Technologies, Inc, Coralville, IA) and sequencing libraries were prepared using the Nextera IDT kit (Illumina, San Diego, CA). The resulting pooled amplicon library was sequenced on an Illumina NovaSeq instrument using paired-end chemistry (2×250bp) at the University of Colorado Anschutz Medical Campus’ Genomics and Microarray Core. Demultiplexed paired-end reads generated from sequencing were imported in QIIME2 version 2020.11^41^. Amplicon sequence variants (ASVs) were generated using DADA2^42^, which was also used to filter reads for quality, remove chimeric sequences, and merge overlapping paired-end reads. Forward and reverse reads were truncated at 248 bp and 250 bp, respectively. Taxonomy was assigned using a Naïve Bayes classifier trained on the Greengenes version 13_8 99% OTUs database^43^, where sequences had been trimmed to include only the base pairs from the V3-V4 region bound by the 341F/805R primer pair.

### Optimization samples

From the study samples, five were selected based on their relatively high amount of *Mh* based on 16S sequencing. As described above, five technical replicates were processed for each sample to test five different concentrations of baits and processing methods. To obtain sufficient amounts of fragmented DNA for this study, multiple aliquots of the same extracted DNA were acoustically sheared with the Covaris M220, cleaned with SPRI beads as described in the supplemental protocol, and combined for TapeStation analysis. From these combined DNA samples, duplicate 250ng aliquots from each of the 5 optimization samples were then used as input for 4 protocol modifications (FULL, HALF, DOUBLE HALF, and DOUBLE QUARTER) and 4 samples for the QUARTER protocol ([5 optimization samples × 2 technical replicates × 4 protocol modifications] + [4 samples × 2 technical replicates × 1 protocol] = 48 total sequencing library preparation attempts). Only one attempt was made at library preparation per replicate. For each of the 5 protocol modifications, 1-2 replicates failed to produce final libraries with DNA concentrations ≥1ng/μl and therefore were not sequenced or included in downstream analysis. This included 1 FULL, 2 HALF, 4 QUARTER, 2 DOUBLE HALF, and 2 DOUBLE QUARTER replicate libraries. The remaining 39 sequencing libraries were pooled in equimolar amounts and sequenced on a single S4 lane on an Illumina NovaSeq 6000 instrument using paired-end chemistry (2×150bp) at the Texas A&M University Institute for Genome Sciences and Society (College Station, TX), with an expected mean sequencing depth of 62.5M paired-end reads per sample.

Sequencing reads were processed with QC trimming, read merging, read deduplication, host DNA removal, and taxonomic classification with Kraken2 and the core NT database. The number of reads classified as *Mh* (on-target) were then compared to identify the library preparation protocol that produces the best results.

### Bioinformatic workflow development

#### Taxonomic classification of Mannheimia haemolytica

Many factors affect the performance of a bioinformatic workflow, starting with sequence quality control, host removal, and taxonomic classification. QC read trimming was performed using Trimmomatic^21^, and host sequences were removed following alignment to a host genome with bwa^24^. Additionally, we identified that our TE library preparation produces DNA fragments with an average length of 250nt, meaning that our 150nt-long sequencing reads could be merged and increase our average read lengths. Therefore, we included a step to merge and deduplicate our reads with FLASH and seqKit^22,23^. Merged and unmerged reads would then be analyzed in parallel, with results combined per sample at the end of the workflow.

For classification, we tested using a simple alignment protocol with bwa to a database of all *Mh* GenBank reference genomes, however, we identified that reads often had multiple alignments and by re-classifying those reads with the coreNT database, we discovered many false positive classifications to other taxa in the same taxonomic lineage as *Mh* (data not shown). We then tested Kraken2, which can handle large databases and employs k-mer scoring to classify reads to the lowest common ancestor (LCA). We tested various pre-made Kraken2 database sizes available on Beg Langmead’s GitHub (https://benlangmead.github.io/aws-indexes/k2). This included the standard database (refSeq archaea, bacteria, viral, plasmid, human, and UniVec_Core) as well as the PlusPF (Standard plus Refeq protozoa, fungi & plant) and the core_NT database (which includes all of RefSeq, TPA, and PDB). We also tested their “minimized” versions, which are made by down sampling the k-mer database to decrease memory requirements. Based on our results, we found that the more comprehensive the database, the more reads we can classify at the species level and reduce false-positive calls, especially compared to minimized databases (data not shown). Going forward, we recommend using the core NT database or a similarly comprehensive database for species-level classification.

#### Variant-level classification

Taxonomic classification past species, to the “strain” or “variant” level, can be challenging, particularly as few genomes are characterized past the strain level. Therefore, we created our own variant-level taxonomic labels based on genome similarity between all *Mh* genomes in our database (n = 2,435). Hierarchal clustering was performed based on the pairwise ANI results using the hclust function with default settings in R (version 4.4.2). We then used cutree to create GSV annotations starting with just 2 groups, and increasing the group number sequentially, up to a total of 30 unique GSV groups (Supplementary File 6). To determine the optimal classification structure for GSVs while minimizing false-positive annotations, we developed custom Python scripts to generate simulated samples containing sequences from varying numbers of different GSVs and test their classification performance as the number of unique GSVs in the database increases- from 2 to 30 GSVs per set (detailed explanation here: https://github.com/Microbial-Ecology-Group/VARIANTplusplus/blob/master/docs/Benchmarking_explanation.md). For each set of GSV annotations, we tested the inclusion of simulated reads from an increasing number of GSVs (i.e. 0, 1, 3, 5, 7, 9) in each simulated sample. Then, for each GSV randomly selected to be represented in a sample, 5 genomes were randomly selected and concatenated to make a single file with 5 genomes per GSV. The whole-genome sequences were then processed with InSilicoSeq^44^ to create 150nt-long reads based on a NovaSeq sequencing error profile and a mean DNA fragment size of 311, at a randomly selected sequencing depth (250000, 500000, 1000000, 2000000). Reads were then merged using FLASH and the resulting merged and unmerged reads were analyzed in parallel. Taxonomic classification was performed with Kraken2 and the core NT database, iterating over confidence values (0, 0.1, 0.2, 0.3, 0.4, 0.5). Reads classified as *Mh* lineage at the species level were extracted with KrakenTools^45^ and aligned to our *Mh* database with Themisto. The alignments were then analyzed with mSWEEP (Metagenomic Sequence Weighted Estimation of Proportions), which applies a Bayesian model to estimate the relative abundance of closely related sequences within the reference database based on the provided grouping file (GSV annotation sets 2-30).

Classification results from the mock samples were parsed to calculate precision, defined as the number of correctly identified GSV groups (true positives) divided by the total number of identified GSV groups (true positives plus false positives), and recall, defined as the number of correctly identified GSV groups divided by the total number of expected GSV groups (true positives plus false negatives). One thousand iterations were conducted for each number of unique GSVs in a mock sample, with precision and recall for each annotation set averaged across all iterations and samples containing varying numbers of GSV groups (i.e., 1, 3, 6, 9). For samples with 0 GSVs, off-target genomes from other bacteria in the same Pasteurellaceae family (n= 5852) were processed in parallel to measure false-positive classification rates, allowing us to pinpoint the best combination of: number of GSV clusters, Kraken2 classifier confidence thresholds, and count filtration to account for false-positive classification.

The selected GSV classification annotation set was then further characterized by plotting a tree using iTOL^46^ (v7) with all reference *Mh* genomes and adding GSV labels in addition to genotype labels and comparing genome membership. Trees created in iTOL were combined into a single figure created with BioRender.com(Figure 2). To assess homology within and between GSV groups, the average ANI was calculated for all genomes within each group. Then, representative genomes for each GSV were selected using dRep’s ‘dereplicate’ workflow, and pairwise ANI distances were then calculated between these representatives. *The VARIANT++ bioinformatic workflow*: We developed the VARIANT++ bioinformatic pipeline using Nextflow DSL2 programming language to replicate our proposed workflow. The pipeline is initiated by utilizing fastQC^47^ and multiQC^48^ to create HTML sequencing quality reports. It then performs quality control trimming of raw sequencing samples using Trimmomatic ^14^, followed by merging of reads when possible. From here, both merged and unmerged reads are processed in parallel, starting with deduplication using seqKit to account for PCR bias resulting in duplicate reads. Then, samples undergo removal of host DNA with bwa and samtools. Non-host reads then undergo taxonomic classification using Kraken2 and the core NT database, as described above with a kraken confidence value of 0.1. After classification, reads belonging to a user-selected taxonomic rank (based on simulated samples, we recommend species) in the lineage of interest (*Mh* in this study) are extracted from the kraken2 output using the ‘extract_kraken_reads’ function with the ‘--include-children’ flag and using the ‘--taxid’ flag from the KrakenTools suite of scripts ^45^. These filtered reads are then aligned to the custom database described above (representing *Mh* in this study) using Themisto, and subsequently classified using mSWEEP and a corresponding GSV annotation set.

A custom python script in the VARIANT++ pipeline then parses the mSWEEP results into count matrices containing the count of reads classified to each GSVs (here, 8 unique GSV groups). Based on the benchmarking results showing the presence of false-positive GSV classifications, even if reads originating from off-target genomes, we elected to apply a count filter to each sample, where the estimated number of false-positive reads from each GSV is calculated and subtracted from the count totals.

Users can execute the complete workflow, from raw reads to count matrix, by including the “--pipeline “full_GSV_pipeline” flag or run specific components individually, such as quality filtering and read merging, deduplication, host removal, taxonomic classification with read extraction, and GSV classification. Importantly, the GSV annotation scheme and VARIANT++ pipeline can be applied to any lineage of interest with a simple substitution of the database and underlying whole-genome comparisons. Instructions and code to replicate the workflow for these data, or to create GSV annotation sets for analysis of other lineages of interest are provided at our GitHub repository (https://github.com/Microbial-Ecology-Group/VARIANTplusplus).

### Investigation samples

*Mh*-TE sequencing libraries were prepared as described using the DOUBLE HALF protocol, as this was determined to be an optimal approach (see above). Library preparation for 1 sample (LOW group) yielded <1ng/μl of DNA and was, therefore, not sequenced and was dropped from downstream analysis. The remaining sequencing libraries (n=19) were pooled in equimolar amounts and sequenced on a single S4 lane on an Illumina NovaSeq 6000 instrument using paired-end chemistry (2×150bp) at the North Texas Genome Center (Arlington, TX), with an expected mean sequencing depth of 131.5M paired-end reads per sample. The raw sequencing reads generated for the Investigation Sample Set using shotgun and TE shotgun sequencing were analyzed using VARIANT++ with the core NT Kraken2 database and the custom *Mh* GSV database as described, providing a count matrix of *Mh* GSVs. These counts were then processed using the R computing environment and packages including phyloseq^49^, vegan^50^, and ggplot2^51^. The full list of R libraries used, count matrices, and R code can be found in our GitHub repository (https://github.com/Microbial-Ecology-Group/Manuscript-Mh_TE_validation). The percent of reads classified to the *Mannheimia* genus and *Mh* species was calculated, as was the percent of counts classified to GSVs. To reduce the potential impact of false-positive classifications, GSV counts were filtered using the calculated false positive rate for each GSV as described above. The comparisons of interest were between samples with different levels of *Mh* abundance (i.e., ZERO, LOW, MEDIUM, HIGH, HIGHEST) and BRD status.

Beta diversity was measured using the vegan R package and the Jaccard dissimilarity index. Pairwise differences in beta diversity were tested using PERMANOVA as employed by the “pairwise.adonis()” R function ^52^. The Benjamini-Hochberg correction was applied to account for multiple comparisons. The dispersion of variance between sample groups was also calculated using the “betadisper” function and tested with the “permutest” function. GSV composition was visualized in heatmap by first adding a small pseudocount of 1 for all taxa, and log transforming counts. Samples in the heatmap (rows) are clustered by *Mh* strata and also labelled to indicate BRD status in addition to a barplot showing the percent of raw reads classified as *Mh* for each sample.

### Shotgun sequencing

#### Shotgun sequencing of Optimization and Investigation Samples

All samples that underwent *Mh* TE sequencing were also sequenced in parallel using traditional shotgun sequencing (n=5 optimization samples, and n=19 investigation samples). One optimization sample was randomly selected to be included in the investigation sample set, and therefore 23 samples were prepared for shotgun sequencing. Briefly, sequencing libraries were prepared using the Illumina DNA Prep Kit following manufacturer’s instructions, using 100ng of DNA for input. Resulting libraries were pooled in equimolar amounts and sequenced on an Illumina NovaSeq 6000 instrument using paired-end chemistry (2×150bp) at the North Texas Genome Center (Arlington, TX) with an expected mean sequencing depth of 109M paired-end reads per sample.

## Data availability

All sequence reads are available through BioProject PRJNA1309097 at the NCBI’s Sequence Read Archive. The code and instructions for the bioinformatic and statistical analyses can be found at the GitHub repository (https://github.com/Microbial-Ecology-Group/Manuscript-Mh_TE_validation) and by the corresponding DOI: 10.5281/zenodo.16989755.

## Author contribution statement

PSM conceptualized the research and directed all aspects of the study, including the development of laboratory and computational methodology, and provided oversight of laboratory and computational analysis, interpretation of results, and oversight of manuscript writing and editing. ED contributed to overall methodology, oversaw software development, data analysis, and data visualization, drafted the original manuscript, and participated in review and editing. LJP contributed to development of laboratory and computational methodology, data analysis and interpretation, contributed to drafting of the original manuscript, and participated in review and editing. CAW oversaw all laboratory analyses, including development of laboratory methodology and interpretation of results, and contributed to manuscript review and editing. RVC contributed to study methodology and assisted with review and editing. WBC contributed to sample collection and to manuscript review and editing. MLC contributed to methodological development and manuscript review and editing. ARW contributed to study conceptualization, methodological development, and manuscript review and editing. NRN contributed to development of computational methodology and manuscript review and editing.

## Notes

https://github.com/Microbial-Ecology-Group/VARIANTplusplus

### Competing Interest Statement

The authors have declared no competing interest.

