## Supplementary figures and images for "Target-enriched sequencing enables genomic characterization within diverse microbial populations – a preprint"

### Supplementary Fig 2

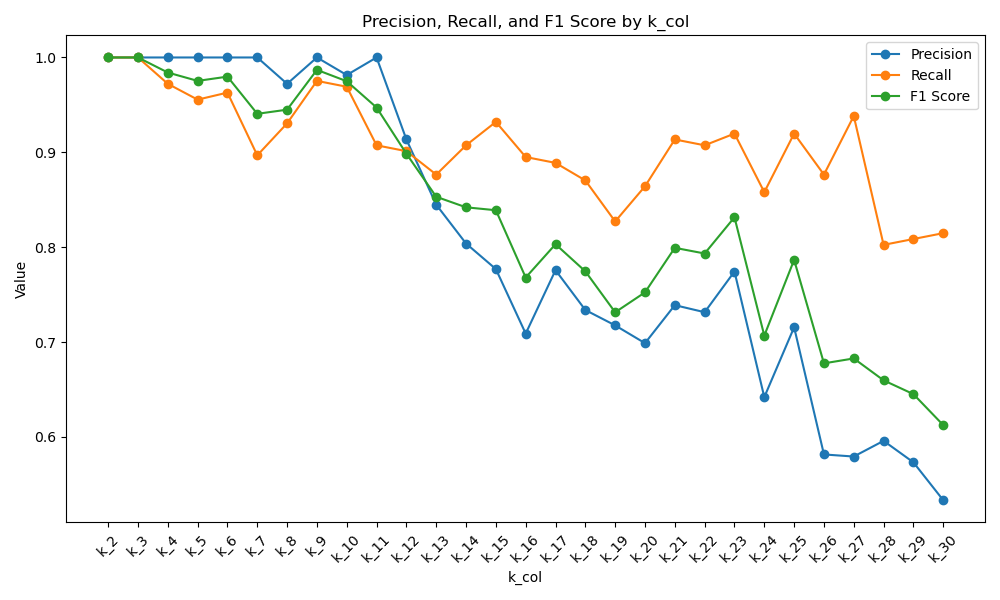

### Supplementary Fig. 1

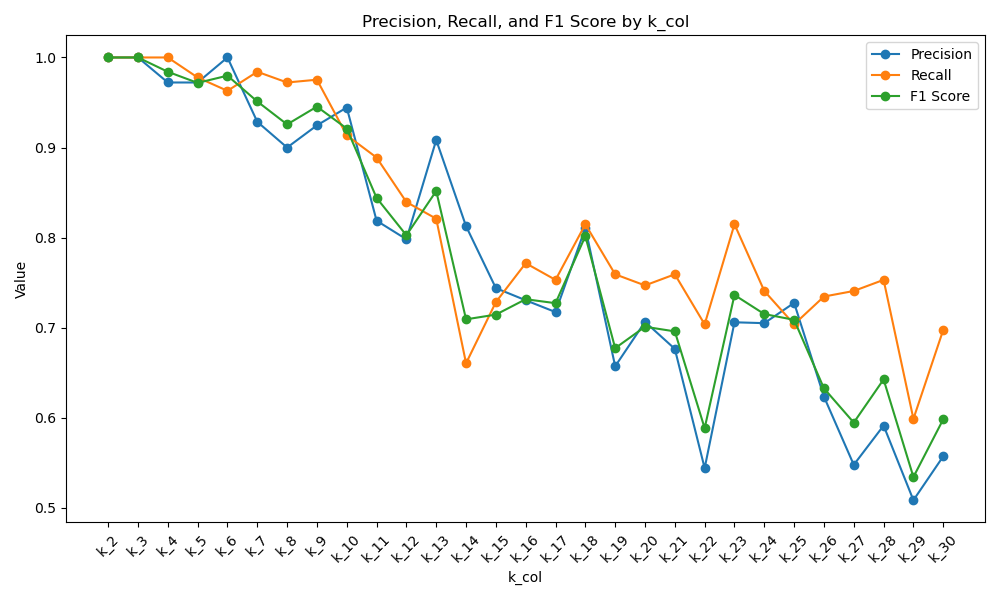

### Supplementary Fig. 3

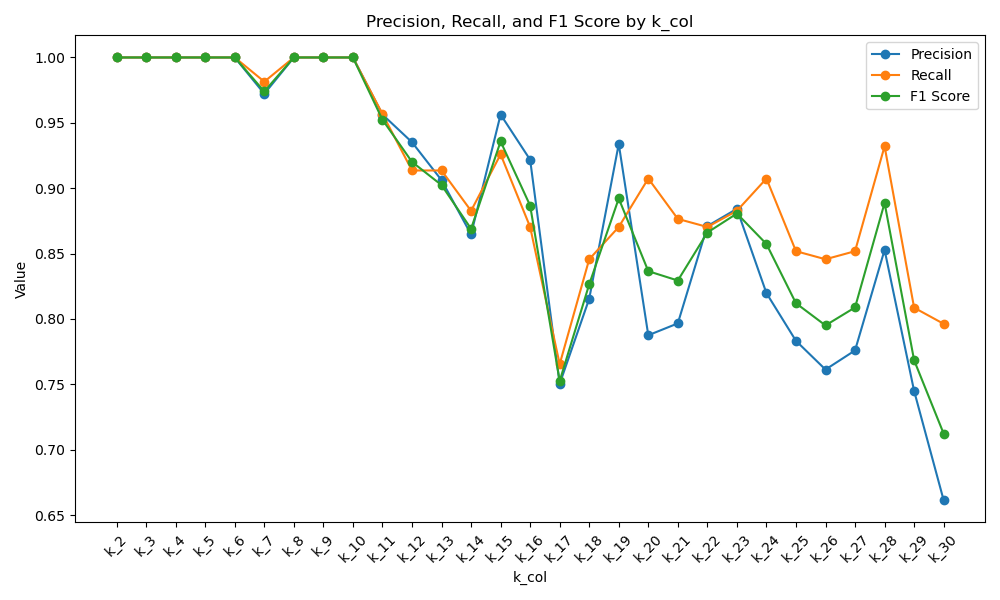

### Supplementary Fig. 4

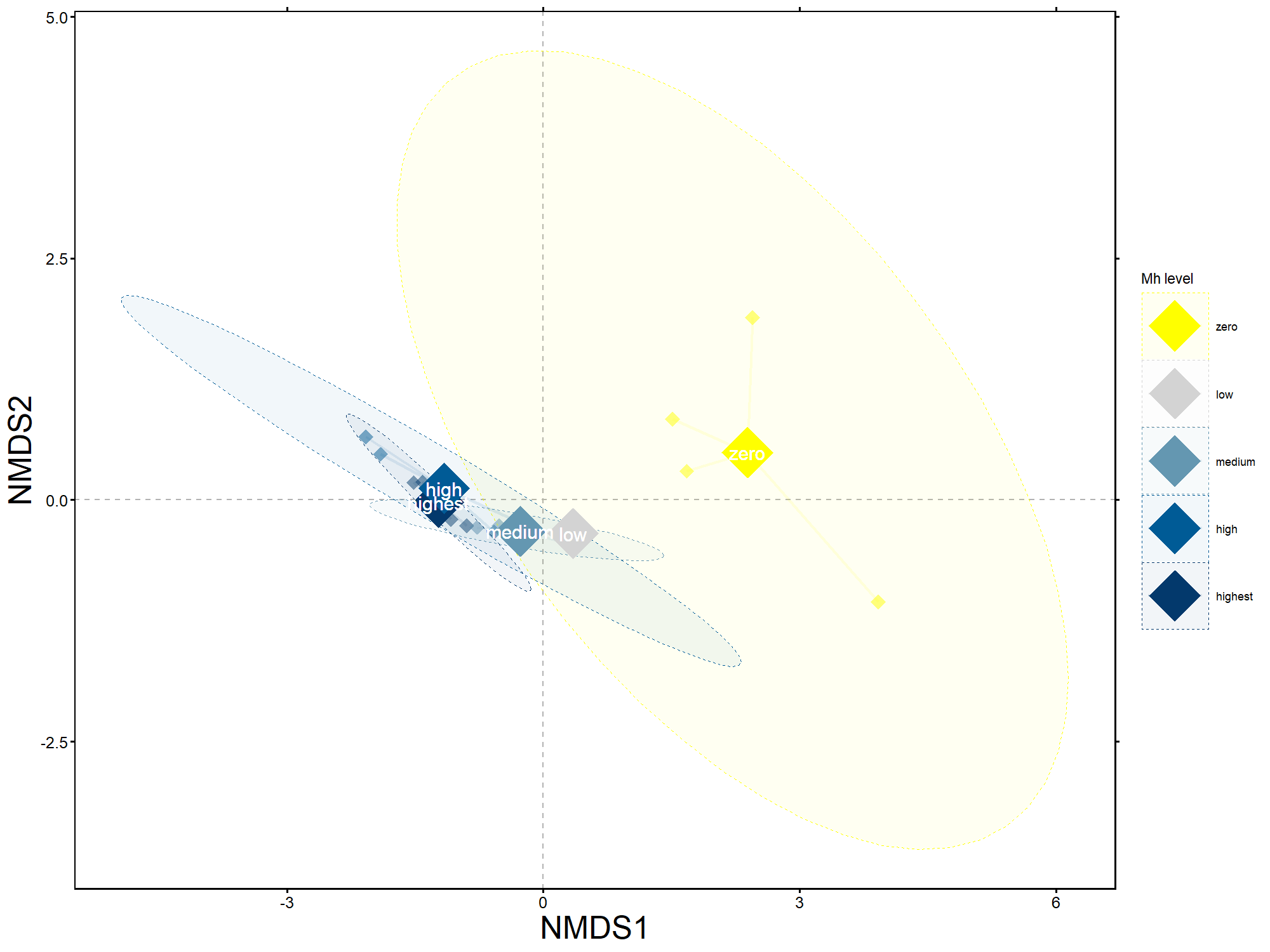
